# Variations in phenolic levels in grapevine buds at eco-dormancy after chemically-induced stress conditions

**DOI:** 10.1101/2021.03.19.436201

**Authors:** Ioannis Daskalakis, Maritina Stavrakaki, Kyriaki Sotirakoglou, Katerina Biniari

## Abstract

Viticulture is one of the briskest economic activities worldwide. A major obstacle impeding such grape yields to satisfy the demand for increased production is the insufficient period of chilling temperatures which, due to climate change, is becoming briefer. The shorter period of cold leads to poor budbreak which, in turn, results in reduced yields. To combat this issue, agronomists have resorted to treatments with chemical regulators to meet the chilling requirement for bud dormancy release several fruit-bearing plants have, including grapevines.

This study aimed at identifying and quantifying during eco-dormancy the individual polyphenolic compounds, and their possible variations, in the latent buds of the ‘Prime^©^’ and ‘Ralli’ table grape cultivars. The study induced chemical stress by means of four different solutions, at three discrete dates per year, for three consecutive years. Phenolics in the latent buds of the studied varieties were analyzed via HPLC. Their quantitative analysis showed variations both between the varieties and between the samples of those varieties collected after the eight treatments. The analysis indicated that the concentration of phenolics continued progressing during the annual growth cycle of the grapevine, with strong positive correlations being detected between catechin, epicatechin, chlorogenic acid, o-coumaric, piceid, procyanidins B1-B2, rutin, ε-viniferin. Luteolin registered a positive correlation with quercetin, resveratrol, o-coumaric, but not with the remaining polyphenols. The results confirmed that (a) the concentration of phenolics of the latent buds is affected by chemically-induced stress conditions, (b) depending on the date of application, significant changes appear in the variations of those phenolics.

**One sentence summary:** Changes in the levels of polyphenol compounds of the grapevine’s latent buds hints, for the first time, how much stress the buds actually undergo under chemically-induced stress conditions

## Introduction

Polyphenols, a group of substances with a broad spectrum of physiological activities, are widespread in plants and are also known for their use in traditional medicine and contemporary medical systems. Polyphenolic acids and their derivatives are carbon-based compounds and have been shown to play a role in tissue browning, flavor, and color characteristics of the derived products (Ho et al., 1992). An understanding of polyphenolic composition in fresh fruit; and of the factors that affect polyphenolic compounds are both critical in the design of products and their storage conditions (Oleszek et al., 1994). Consequently, many have been the studies which undertook in the past to investigate the polyphenolic acid compositions of various fruits -and their related cultivars, such as apples and pears (Blankenship et al., 1985; Amiot et al., 1993); *Pyrus* (Challice et. al., 1968); other pome and stone fruits (Herrmann, 1989); *Diospyros* (Ayaz et al., 1997); carrots (Babic et al., 1993); and *Prunus* (Nortje and Koeppen, 1965).

Polyphenolic compounds are active biological molecules having one or more benzene rings with one or more hydroxyl functions (Manach et al., 2004). At plant level, polyphenolic compounds contribute to development, cell multiplication, reproduction, differentiation, flowering, and lignification of a plant. As to their concentration in plants, that depends on a number of genetic, physiological and environmental parameters (Balasundram et al., 2007). Polyphenolic compounds accumulate mainly in the cell membrane (lignin and some flavonoids) and in vacuoles where soluble polyphenols such as chlorogenic acid, anthocyanins, flavonols, and tannins are stocked (Peer et al., 2001). At the same time, substances such as polyphenolic compounds, amino acids (namely prolin), and the group of polyamins (PAs) are known to be involved in the resistance response to stress by many plant species, the grapevine included. What is more, they act as free radical scavengers (Bouchereau et al., 1999)

The grapevine follows a growth cycle that consists of several steps (Pouget, 1963). The most important of those steps is dormancy, whether it characterizes the dormancy of the latent buds (endo-dormancy) triggered endogenously; or the dormancy of the vine (eco-dormancy). In the vineyard, eco-dormancy is triggered by a sharp drop in temperature. When cold and low temperatures give rise to challenging environmental conditions for vines during their annual cycle, plant growth slows down, followed by total arrest in autumn and winter. The vines then become dormant, a particular physiological condition that allows them to withstand climate adversities. However, regarding the installation of bud dormancy that takes place earlier during the growth cycle, the decreases in the photoperiod are considered one of the most significant factors among those that can induce that installation (Febvre, 1981; Olsen, 2006; Daskalakis and Biniari, 2019).

The removal of dormancy is also controlled by environmental conditions. When it comes to grapevines, the latent buds acquire the ability to clog under the effect of low temperatures (0 °C - 10 °C). Thus, cold in temperate climates remains the most effective way to break dormancy in woody species (Nigond, 1967; Champagnat, 1983).

Due to the climate change, the current duration of chilling temperatures requisite for bud dormancy release of several fruit plants, grapevines included, suffices no longer. Worse, the brevity of that requisite chilling period appears to be intensifying. As a result, growers and agronomists have been increasingly resorting to treatments of such fruit plants with chemical and growth regulators. Examples include chemicals, used commercially by many wine-producing countries, such as mineral oil, dinitro-O-cresol (DNOC), thiourea, calcium cyanamide, potassium cyanamide, hydrogen cyanamide, and garlic paste (Iwasaki, 1980; Zelleke and Kliewer, 1989; Kubota and Miyamuki, 1992; Mizutani et al., 1994; Botelho et al., 2007). Additionally, calcium cyanamide and hydrogen cyanamide have been documented as being quite effective on grape bud dormancy release (Weaver et al., 1961; Iwasaki, 1980; Shulman et al., 1983; Kuroi, 1985; Lin and Wang, 1985). Hydrogen cyanamide is another chemical that has been used successfully to supplement the chilling requirement, significantly improving budbreak (Dokoozlian et al., 1995; Carreno et al., 1999).

To try to comprehend this resting phase of the grapevine, several authors have studied the main biochemical changes the latent buds undergo. In this phenomenon of dormancy, some researchers have made special mention of the involvement of hydrogen peroxide (H_2_O_2_) (Lavee and May, 1997), and polyphenolic compounds (Nagar and Sood, 1996; Macheix et al., 2005; Zohra et al., 2011). The evolution of levels in polyphenolic compounds of latent buds during their annual growth cycle seems to be related to that of hydrogen peroxide (H_2_O_2_). This hypothesis is supported by Li et al. (2011) who reported that the cold treatment leads to higher CAT (catalase) and APX (ascorbate peroxidase) activities in the leaves, since polyphenolic compounds are essentially an anti-stress mechanism.

However, so far, no studies have been carried out on the individual polyphenolic compounds in the latent buds of the grapevine during eco-dormancy and, more importantly, under chemically-induced stress conditions.

Thus, the purpose of the present study was (a) to examine the premise that oxidative stress is caused by applications with hydrogen cyanamide which advance budbreak; and (b) to monitor, for the first time during such experiments and during the annual vegetative cycle, the variations in the levels of thirteen (13) individual polyphenolic compounds of the latent buds in two (2) grapevine varieties ‘Prime^©^’ and ‘Ralli’ *(Vitis vinifera* L.), also in relation to the different time of applications and the two different chemical substances used (Theocopper & Erger), for three consecutive years.

## Materials and Methods

### Plant material and experimental design

‘Prime^©^’ is a white table grape variety and ‘Ralli’ is a red table grape variety *(Vitis vinifera* L.). Both are regarded by viticulturists worldwide as two of the grapevine varieties with the earliest maturation. The present study’s researchers located seven-year old vines of these two varieties in a vineyard in Corinth (alt: 10 m, gradient: 2%), northeastern Peloponnese, Greece. The selected vines (already grafted on rootstock 1103 Paulsen), were bilaterally cordon-trained (bilateral Royat) at 2.2 m x 1.2 m intervals, and cane-pruned to 10-node canes per arm. Each vine consisted of four (4) arms and, therefore, each vine comprised four (4) canes in total. The usual viticultural techniques entailed: fertilization using 11-15-15 NPK at a dose of 250 g/vine; canopy management techniques (shoot thinning, topping; girdling); and irrigation. All studied vines had been grown in the same area and under the same climate and soil conditions.

### Treatments

The experiment lasted three (3) consecutive years and, more specifically, during the 2016-2018 cultivation seasons. Four (4) different solutions (Theocopper & Theocal, Dormex, Erger, and garlic extract), known for their ability to advance budbreak, were applied to and evaluated for two table grape varieties, ‘Prime^©^’ and ‘Ralli’. Eight (8) preparations were applied to each variety at three different dates (December 15, January 15, and February 15), in three (3) consecutive years, bringing the total of treatments to one hundred and forty-four (144). In other words, the two (2) varieties were treated to eight (8) applications, on three different (3) dates, for three (3) consecutive years (2×8×3×3=144). The groups of ten (10) vine canes selected were chosen not only because of their morphology but also because they were the most representative of each variety. For the needs of the experiment, the research made use of a Randomized Complete Block Design with three (3) replications per treatment. Each group of 10 vine canes constituted one (1) replication.

The eight applications were effected using the following mixtures:

i. 10 mL Theocopper and 1 g Theocal per Liter [TA];
ii. 20 mL Theocopper and 1 g Theocal per Liter [TB];
iii. 25 mL Dormex^®^ per Liter [DA];
iv. 50 mL Dormex^®^ per Liter [DB];
v. 35 mL Erger and 80 mL Active Erger per Liter [EA];
vi. 70 mL Erger and 160 mL Active Erger per Liter [EB];
vii. extract from 300 g (FW-fresh weight) garlic evaporated and then diluted in 1 L water [GA]; and
viii. distilled water (control treatment) [Control].

Preparations were applied using a knapsack sprayer. The synthesis of the preparations was as follows: Theocopper: 10% sugars, 10% amino acids, 12% urea, 1.4% potassium in organic form, 12% nitrogen in organic form, and 3.5% organic matter; Theocal: 30% calcium, 35% organic matter with a pH of 7.1. Dormex^®^: H_2_CN_2_ 490 g L^-1^ (Basf Co.)

As to the garlic extract, that was prepared as follows: 300 g of fresh and peeled cloves of garlic *(Allium sativum* L.) were ground in a blender. The ensuing mash was blended again with 1 L distilled water, filtered, and raised to 2 L by the addition of distilled water to obtain a final concentration of 15% garlic extract (Kubota and Miyamuki, 1992).

### Sampling

The canes were sampled on day 0, i.e., before the first application, and then on day 3, 6, 9, 15, and 30 following each application. As stated earlier, the applications took place on three different dates (December 15; January 15; February 15) with the sprayings focusing on the first ten (10) latent buds starting at the base of the canes. The samples were randomly collected from the entire vineyard so that sampling may be more homogeneous and representative of the space the samples had been grown in. Once collected, they were frozen in liquid nitrogen and then lyophilized. Last, buds were homogenized in liquid nitrogen using a pre-chilled mortar and pestle for the extraction.

### Reagents and chemicals

The various polyphenolic compounds analyzed were identified according to their order of elution and the retention times of the pure compounds. Non-colored polyphenolics were purchased from a number of different sources. More specifically, catechin, vanillic acid, chlorogenic acid, epicatechin, o-coumaric acid, and rutin were procured from Sigma, St. Louis, MO, USA. Luteolin, procyanidin B1, procyanidin B2, ε-viniferin, quercetin, trans-resveratrol, and piceid were purchased from Extrasynthese, Gemay, France.

### Extraction of polyphenols

The procedure followed in order to extract the polyphenols from each and every sample was the following: 5 mL of 70% v/v methanol acidified with 1% formic acid (v/v) were added to 200 mg of dried tissue which were then weighed on a precision scale (KERN 410). The mixture was shaken and placed in a water bath for 60 min at a temperature of 40 °C and centrifuged at 4,000 rpm for 6 min. Once the supernatants (5 mL) were collected, the same extraction process was repeated twice more for the remaining polyphenols, resulting in a total extraction volume of 15 mL 70% v/v methanol. All fractions were combined. Last, the supernatants were stored at a temperature below 80 °C until the time of the analysis.

### Analysis by HPLC

In order to measure individual polyphenols with HPLC, liquid extraction was carried out: 2 mL of the 70% v/v methanol extract were evaporated with a sample concentrator at room temperature under a stream of nitrogen gas. By means of that method, the tubes held nothing but the aqueous residue containing the polyphenolic compounds. Next, 1 mL water (HPLC grade) was added to that residue, followed by vortex. Then, 2 mL of ethyl acetate were added to the mixture followed by vortex for 1 min. The following step entailed transferring the supernatant ethyl acetate to a new tube, adding 2 mL of ethyl acetate to the pellet, followed by vortex for 1 min. The new supernatant ethyl acetate was transferred and mixed with the previous one. The combined ethyl acetate supernatant was evaporated with a sample concentrator at room temperature under a stream of nitrogen gas and the pellet was dissolved in 1 mL of methanol (HPLC grade). Last, prior to the HPLC analysis, the extract was filtered through a 0.22 μm membrane.

Monomeric and dimeric phenols [(+)-catechin, (-)-epicatechin, procyanidins B1 and B2, vanillic acid, o-coumaric, chlorogenic acid, piceid, rutin, trans-resveratrol, ε- viniferin, quercetin, and luteolin] were determined by the HPLC-DAD system (Shimadzu Nexera).

For the separation of monomeric and dimeric polyphenols, the study employed a 250 x 4.6 mm ID, 5 μm, Waters x select C18 column operating at 25°C. The eluent comprised (a) H_2_O/ C_2_H_4_O_2_ (99.3:0.7); and (b) CH_3_OH (100). The flow rate stood at 0.5 mL min^-1^. The linear gradient program used for the elution was the one described in Biniari et al. (2020).

### Data analysis

The experiment was designed and implemented following the principles of the Randomized Complete Block Design. Data were analyzed using the Statgraphics Centurion statistical package (Statgraphics Technologies, Inc., version 17) and are presented as mean ± SE (Standard Error) of the three (3) replications. Each group of ten (10) grapevine canes constituted one (1) replication.

A repeated measures General Linear Model (GLM) was applied to the data (polyphenolic compounds) considering the sampling time as repeated measure, with fixed effects of the treatment (control, TA, TB, DA, DB, EA, EB, GA), the grapevine variety (Prime^©^, Ralli), the year of the experiment (2016, 2017, 2018), the dates when the applications were performed (December 15, January 15, and February 15) and number of days that had elapsed between applications and sampling (0, 3, 6, 9, 15, 30 days). *Post hoc* analyses were performed using Tukey’s HSD test. The Kolmogorov-Smirnov test revealed that all variables followed normal distribution. Moreover, in order to reduce the dimensionality of the data and investigate the relationships between and among polyphenolic compounds, pooled data (all polyphenols) were subjected to principal component analysis (PCA). Results and their detailed description are shown in the present study’s tables and diagram (plot). The significance level for all tests was set at 5%

## Results and Discussion

According to their respective retention time, thirteen (13) polyphenolic compounds were identified: (1) procyanidin B1; (2) procyanidin B2; (3) catechin; (4) chlorogenic acid; (5) vanillic acid; (6) epicatechin; (7) piceid; (8) rutin; (9) o-coumaric; (10) resveratrol; (11) ε-viniferin; (12) quercetin; and (13) luteolin. The polyphenolic content is expressed as μg equivalent per g of dry weight (μg g^-1^ dw).

The content of polyphenols in ‘Prime^©^’ and ‘Ralli’, in tandem with (i) the year of the experiment; and (ii) the month when the applications were performed, is presented in Table 1. The content of all individual polyphenolic compounds in the two varieties under study registered their highest point in the application effected in February, with the exception of vanillic acid and epicatechin whose content peaked in January (Table 1). With regard to the year when the experiment took place, the results showed that polyphenolics procyanidin B1, catechin, vanillic acid, epicatechin, piceid, rutin, and ε-viniferin exhibited their highest concentration in 2016, the first year of the experiment (Table 1).

**Table 1:** Variation of individual polyphenolic compounds in the buds at eco-dormancy in relation to the year and month the applications were carried out

|  | Year |  |  |  |  | Month |  |  |  |  |
| --- | --- | --- | --- | --- | --- | --- | --- | --- | --- | --- |
|  | 2016 | 2017 | 2018 | SEM | P-Value | December | January | February | SEM | P-Value |
| **Procyanidin B1** | 1284.5<sup>c</sup> | 850.9<sup>a</sup> | 970.8<sup>b</sup> | 7.21 | <0.001 | 955.3<sup>a</sup> | 1032.1<sup>b</sup> | 1118.6<sup>c</sup> | 7.15 | <0.001 |
| **Procyanidin B2** | 877.1<sup>a</sup> | 966.6<sup>b</sup> | 961.9<sup>b</sup> | 8.11 | <0.001 | 875.7<sup>a</sup> | 898.8<sup>a</sup> | 1031.1<sup>b</sup> | 8.04 | <0.001 |
| **Catechin** | 816.7<sup>c</sup> | 541.6<sup>b</sup> | 384.8<sup>a</sup> | 6.78 | <0.001 | 513.9<sup>a</sup> | 612.1<sup>b</sup> | 617.1<sup>b</sup> | 6.72 | <0.001 |
| **Chlorogenic acid** | 214.3<sup>ab</sup> | 219.1<sup>b</sup> | 211.1<sup>a</sup> | 1.87 | <0.001 | 188.8<sup>a</sup> | 203.8<sup>b</sup> | 251.7<sup>c</sup> | 1.85 | <0.001 |
| **Vanillic acid** | 126.1<sup>c</sup> | 65.66<sup>a</sup> | 77.88<sup>b</sup> | 1.19 | <0.001 | 77.31<sup>a</sup> | 97.69<sup>b</sup> | 94.65<sup>b</sup> | 1.17 | <0.001 |
| **Epicatechin** | 2216.9<sup>c</sup> | 1124.6<sup>a</sup> | 1724.0<sup>b</sup> | 9.85 | <0.001 | 1488.6<sup>a</sup> | 1843.8<sup>b</sup> | 1733.1<sup>c</sup> | 9.77 | <0.001 |
| **Piceid** | 125.3<sup>c</sup> | 99.45<sup>b</sup> | 74.58<sup>a</sup> | 0.88 | <0.001 | 94.24<sup>a</sup> | 97.02<sup>b</sup> | 108.1<sup>c</sup> | 0.88 | <0.001 |
| **Rutin** | 2206.5<sup>c</sup> | 1866.0<sup>b</sup> | 1606.7<sup>a</sup> | 26.92 | <0.001 | 1788.2<sup>a</sup> | 1809.5<sup>a</sup> | 2081.4<sup>b</sup> | 26.7 | <0.001 |
| **o-coumaric** | 160.1<sup>b</sup> | 134.1<sup>a</sup> | 178.5<sup>c</sup> | 2.16 | <0.001 | 130.9<sup>a</sup> | 155.4<sup>b</sup> | 186.4<sup>c</sup> | 2.14 | <0.001 |
| **Resveratrol** | 145.2<sup>a</sup> | 161.1<sup>b</sup> | 161.2<sup>b</sup> | 1.77 | <0.001 | 121.2<sup>a</sup> | 162.7<sup>b</sup> | 183.6<sup>c</sup> | 1.75 | <0.001 |
| **ε-viniferin** | 504.1<sup>c</sup> | 280.9<sup>b</sup> | 213.4<sup>a</sup> | 2.74 | <0.001 | 304.3<sup>a</sup> | 325.2<sup>b</sup> | 369.6<sup>c</sup> | 2.72 | <0.001 |
| **Quercetin** | 181.4<sup>a</sup> | 232.3<sup>b</sup> | 301.1<sup>c</sup> | 2.58 | <0.001 | 229.6<sup>a</sup> | 234.4<sup>a</sup> | 250.7<sup>b</sup> | 2.56 | <0.001 |
| **Luteolin** | 88.64<sup>a</sup> | 112.2<sup>b</sup> | 192.4<sup>c</sup> | 1.36 | <0.001 | 115.6<sup>a</sup> | 134.1<sup>b</sup> | 143.5<sup>c</sup> | 1.34 | <0.001 |

Values are the means of triplicates. Values on the same line carrying a different superscript (a - c) are significantly different at $P < 0.05$ . The results are expressed as $\mu\text{g}$ equivalent per g of dry weight ( $\mu\text{g g}^{-1}$ dw)

The polyphenolic content in ‘Prime^©^’ and ‘Ralli’ as it relates to (i) the studied variety; and (ii) the lapse of days between applications and cane sampling, is presented in Table 2.

**Table 2:** Variation of individual polyphenolic content in relation to the variety under study and the lapse of days between applications and cane sampling

|  | Variety |  |  |  | Sampling after |  |  |  |  |  |  |  |
| --- | --- | --- | --- | --- | --- | --- | --- | --- | --- | --- | --- | --- |
|  | 'Prime<sup>®</sup>' | 'Ralli' | SEM | P-Value | 0 day | 3 days | 6 days | 9 days | 15 days | 30 days | SEM | P-Value |
| **Procyanidin B1** | 1088.8<sup>b</sup> | 981.8<sup>a</sup> | 5.84 | <0.001 | 677.1<sup>a</sup> | 1064.2<sup>b</sup> | 1075.3<sup>b</sup> | 1142.9<sup>c</sup> | 1118.4<sup>c</sup> | 1134.2<sup>c</sup> | 9.91 | <0.001 |
| **Procyanidin B2** | 1019.8<sup>b</sup> | 850.6<sup>a</sup> | 6.56 | <0.001 | 637.8<sup>a</sup> | 982.9<sup>b</sup> | 966.6<sup>b</sup> | 1030.7<sup>c</sup> | 987.9<sup>bc</sup> | 1005.1<sup>bc</sup> | 11.14 | <0.001 |
| **Catechin** | 634.8<sup>b</sup> | 527.3<sup>a</sup> | 5.49 | <0.001 | 378.7<sup>a</sup> | 600.5<sup>c</sup> | 561.1<sup>b</sup> | 666.1<sup>d</sup> | 605.1<sup>c</sup> | 674.7<sup>d</sup> | 9.32 | <0.001 |
| **Chlorogenic acid** | 249.4<sup>b</sup> | 180.1<sup>a</sup> | 1.51 | <0.001 | 148.7<sup>a</sup> | 214.2<sup>b</sup> | 219.1<sup>bc</sup> | 230.6<sup>d</sup> | 228.9<sup>cd</sup> | 247.2<sup>e</sup> | 2.57 | <0.001 |
| **Vanillic acid** | 101.1<sup>b</sup> | 78.62<sup>a</sup> | 0.96 | <0.001 | 57.33<sup>a</sup> | 94.73<sup>b</sup> | 93.01<sup>b</sup> | 96.99<sup>b</sup> | 98.5<sup>b</sup> | 98.76<sup>b</sup> | 1.63 | <0.001 |
| **Epicatechin** | 1836.1<sup>b</sup> | 1541.1<sup>a</sup> | 7.98 | <0.001 | 1249.9<sup>a</sup> | 1706.9<sup>b</sup> | 1754.7<sup>bc</sup> | 1768.4<sup>c</sup> | 1739.8<sup>bc</sup> | 1911.3<sup>d</sup> | 13.55 | <0.001 |
| **Piceid** | 109.1<sup>b</sup> | 90.54<sup>a</sup> | 0.72 | <0.001 | 65.91<sup>a</sup> | 100.6<sup>b</sup> | 109.2<sup>c</sup> | 107.9<sup>c</sup> | 106.2<sup>c</sup> | 108.8<sup>c</sup> | 1.22 | <0.001 |
| **Rutin** | 2149.3<sup>b</sup> | 1636.8<sup>a</sup> | 21.81 | <0.001 | 1186.2<sup>a</sup> | 2044.2<sup>bc</sup> | 1967.7<sup>b</sup> | 1965.7<sup>b</sup> | 2188.1<sup>c</sup> | 2006.5<sup>b</sup> | 37.02 | <0.001 |
| **o-coumaric** | 165.8<sup>b</sup> | 149.3<sup>a</sup> | 1.75 | <0.001 | 84.59<sup>a</sup> | 163.3<sup>b</sup> | 170.6<sup>bc</sup> | 174.4<sup>bc</sup> | 173.5<sup>bc</sup> | 178.9<sup>c</sup> | 2.97 | <0.001 |
| **Resveratrol** | 141.3<sup>a</sup> | 170.4<sup>b</sup> | 1.43 | <0.001 | 86.99<sup>a</sup> | 153.1<sup>b</sup> | 163.6<sup>c</sup> | 165.1<sup>c</sup> | 177.9<sup>d</sup> | 188.3<sup>d</sup> | 2.44 | <0.001 |
| **ε-viniferin** | 343.5<sup>b</sup> | 322.6<sup>a</sup> | 2.23 | <0.001 | 209.1<sup>a</sup> | 340.4<sup>b</sup> | 350.6<sup>bc</sup> | 361.2<sup>cd</sup> | 368.6<sup>d</sup> | 368.3<sup>d</sup> | 3.77 | <0.001 |
| **Quercetin** | 248.2<sup>b</sup> | 228.3<sup>a</sup> | 2.09 | <0.001 | 160.9<sup>a</sup> | 247.9<sup>b</sup> | 249.1<sup>b</sup> | 261.2<sup>b</sup> | 255.6<sup>b</sup> | 254.5<sup>b</sup> | 3.55 | <0.001 |
| **Luteolin** | 119.7<sup>a</sup> | 142.4<sup>b</sup> | 1.09 | <0.001 | 90.29<sup>a</sup> | 137.6<sup>bc</sup> | 140.4<sup>cd</sup> | 132.9<sup>b</sup> | 139.3<sup>bcd</sup> | 145.9<sup>d</sup> | 1.86 | <0.001 |

Values are the means ( $\mu\text{g g}^{-1}$ dw) of triplicates. Values on the same line carrying a different superscript (a - d) are significantly different at $P < 0.05$ . The results are expressed as $\mu\text{g}$ equivalent per g of dry weight ( $\mu\text{g g}^{-1}$ dw)

When compared to ‘Ralli’, ‘Prime^©^’ is characterized by a greater concentration in almost all polyphenolics, with the exception of resveratrol and luteolin. Despite the fact that both ‘Prime^©^ and ‘Ralli’ are early maturation varieties, ‘Prime^©^’ sprouts even earlier than ‘Ralli’. Thus, it can be surmised that ‘Prime^©^’ is more sensitive to stress than ‘Ralli’ (Table 2).

The distribution (%) of analyzed polyphenolic compounds within the studied varieties in relation to the lapse of days between applications and cane sampling is presented in Table 2. The results showed that the highest concentrations were observed in the samplings that took place on the 15-day and 30-day mark following the applications.

The total amounts of polyphenolic compounds increase in early July and, by the end of that phase (pre-dormancy), those amounts start decreasing, coinciding with the start of the endo-dormancy phase (Said et al., 2014). During that phase, the amounts of polyphenolic compounds gradually decrease, while the amount of polyphenolic compounds in latent buds dramatically decreases. That rapid decrease occurs within a short period of time that coincides with the end of the endo-dormancy phase when temperatures remain below 10°C for at least seven (7) successive days. Afterwards, their total amounts start increasing slightly until budburst. That moderate increase corresponds to the buds’ dehydration phase (Said et al., 2014).

The polyphenolic content, in relation to the treatments applied to the ‘Prime^©^’ variety, is presented in Table 3, while that of the ‘Ralli’ variety, again in relation to the treatments applied, is presented in Table 4.

**Table 3:**
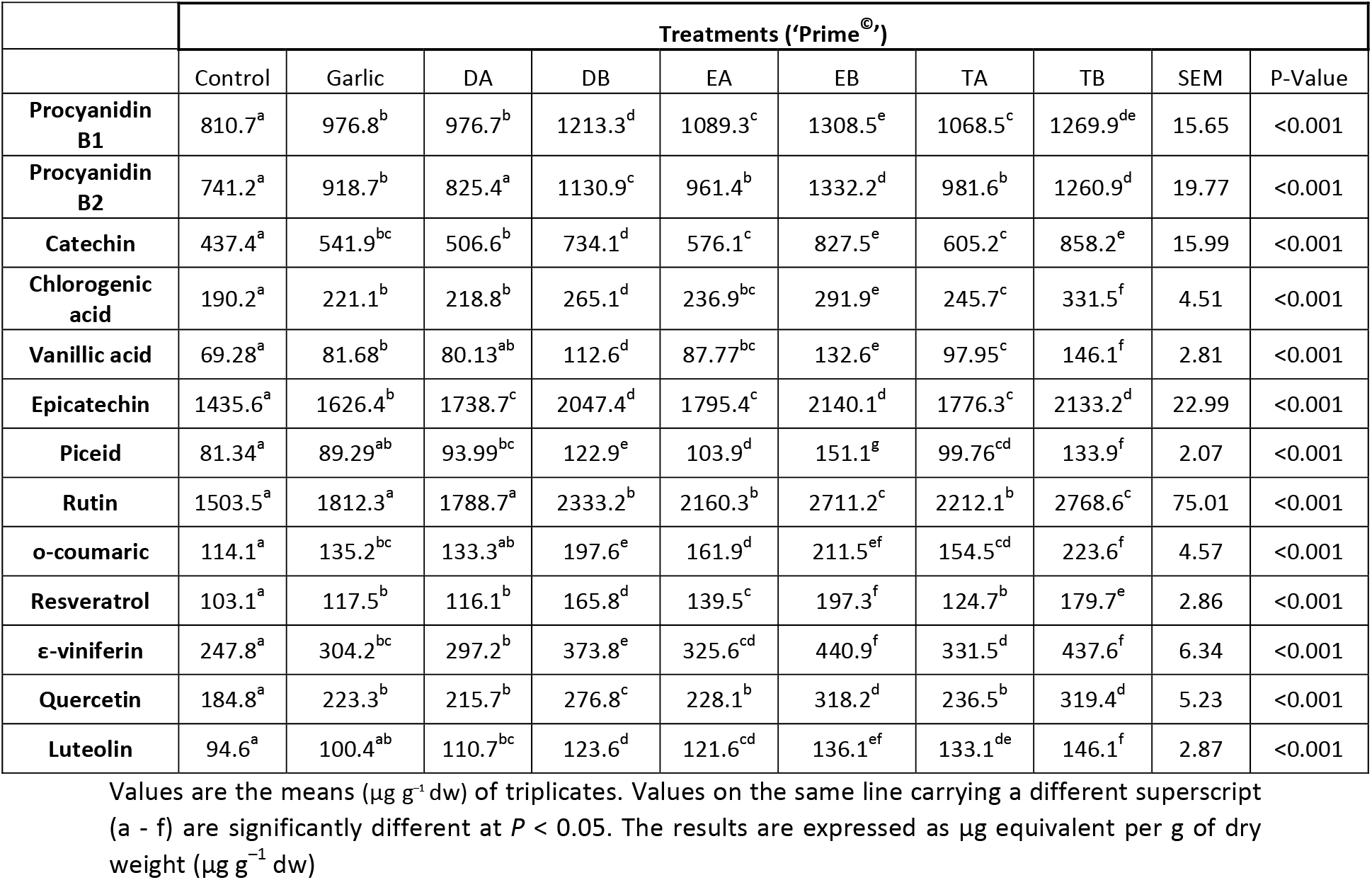
Variation of individual polyphenolic content in relation to the treatment applied on the buds of the ‘Prime^©^’ variety

**Table 4:** Variation of individual polyphenolic content in relation to the treatment applied on the buds of the ‘Ralli’ variety

|  | Treatments ('Ralli') |  |  |  |  |  |  |  |  |  |
| --- | --- | --- | --- | --- | --- | --- | --- | --- | --- | --- |
|  | Control | Garlic | DA | DB | EA | EB | TA | TB | SEM | P-Value |
| **Procyanidin B1** | 737.7<sup>a</sup> | 889.5<sup>bc</sup> | 865.9<sup>b</sup> | 1057.9<sup>e</sup> | 947.9<sup>cd</sup> | 1190.4<sup>f</sup> | 962.3<sup>d</sup> | 1198.6<sup>f</sup> | 15.68 | <0.001 |
| **Procyanidin B2** | 653.1<sup>a</sup> | 752.3<sup>bc</sup> | 701.2<sup>ab</sup> | 955.7<sup>e</sup> | 825.2<sup>d</sup> | 1116.9<sup>f</sup> | 789.3<sup>cd</sup> | 1012.9<sup>e</sup> | 15.48 | <0.001 |
| **Catechin** | 385.2<sup>a</sup> | 445.2<sup>b</sup> | 442.9<sup>b</sup> | 602.7<sup>c</sup> | 488.3<sup>b</sup> | 696.7<sup>d</sup> | 487.6<sup>b</sup> | 675.6<sup>d</sup> | 13.17 | <0.001 |
| **Chlorogenic acid** | 141.2<sup>a</sup> | 155.5<sup>ab</sup> | 158.2<sup>b</sup> | 204.1<sup>d</sup> | 179.4<sup>c</sup> | 231.4<sup>e</sup> | 163.1<sup>b</sup> | 202.2<sup>d</sup> | 3.62 | <0.001 |
| **Vanillic acid** | 56.96<sup>a</sup> | 66.08<sup>abc</sup> | 65.51<sup>ab</sup> | 87.32<sup>d</sup> | 75.39<sup>c</sup> | 106.4<sup>e</sup> | 73.75<sup>bc</sup> | 100.4<sup>e</sup> | 2.22 | <0.001 |
| **Epicatechin** | 1206.5<sup>a</sup> | 1386.8<sup>b</sup> | 1416.7<sup>b</sup> | 1660.3<sup>d</sup> | 1510.9<sup>c</sup> | 1872.1<sup>f</sup> | 1506.4<sup>c</sup> | 1769.6<sup>e</sup> | 19.9 | <0.001 |
| **Piceid** | 67.67<sup>a</sup> | 75.65<sup>b</sup> | 75.21<sup>b</sup> | 103.5<sup>d</sup> | 83.69<sup>c</sup> | 117.8<sup>e</sup> | 83.69<sup>bc</sup> | 117.8<sup>e</sup> | 1.71 | <0.001 |
| **Rutin** | 1160.9<sup>a</sup> | 1334.4<sup>b</sup> | 1405.2<sup>b</sup> | 1763.6<sup>c</sup> | 1436.24<sup>b</sup> | 2057.8<sup>d</sup> | 1642.1<sup>c</sup> | 2149.9<sup>d</sup> | 36.61 | <0.001 |
| **o-coumaric** | 92.93<sup>a</sup> | 115.8<sup>b</sup> | 121.1<sup>bc</sup> | 179.8<sup>d</sup> | 141.1<sup>c</sup> | 195.2<sup>de</sup> | 140.7<sup>c</sup> | 209.2<sup>e</sup> | 4.69 | <0.001 |
| **Resveratrol** | 111.7<sup>a</sup> | 136<sup>b</sup> | 136.1<sup>b</sup> | 189.3<sup>d</sup> | 150.2<sup>bc</sup> | 229.6<sup>e</sup> | 164.2<sup>c</sup> | 237.5<sup>e</sup> | 4.31 | <0.001 |
| **ε-viniferin** | 244.6<sup>a</sup> | 281.2<sup>bc</sup> | 276.1<sup>b</sup> | 345.1<sup>e</sup> | 303.3<sup>cd</sup> | 405.4<sup>f</sup> | 314.2<sup>d</sup> | 404.2<sup>f</sup> | 5.35 | <0.001 |
| **Quercetin** | 169.3<sup>a</sup> | 195.6<sup>b</sup> | 197.6<sup>b</sup> | 246.9<sup>c</sup> | 219.8<sup>b</sup> | 271.9<sup>cd</sup> | 214.5<sup>b</sup> | 293.7<sup>d</sup> | 5.82 | <0.001 |
| **Luteolin** | 114.4<sup>a</sup> | 117.4<sup>a</sup> | 136.5<sup>b</sup> | 147.6<sup>bc</sup> | 143.7<sup>bc</sup> | 155.9<sup>cd</sup> | 151.4<sup>cd</sup> | 162.8<sup>d</sup> | 3.13 | <0.001 |

Values are the means ( $\mu\text{g g}^{-1}\text{ dw}$ ) of triplicates. Values in the same line with different superscript (a - f) are significantly different at $P < 0.05$ . The results are expressed as $\mu\text{g}$ equivalent per g of dry weight ( $\mu\text{g g}^{-1}\text{ dw}$ )

Compared to the control treatment, all other treatments produced an increase in the content of polyphenolic compounds (Tables 3, 4). In all likelihood, the evolution of polyphenolic compounds in the latent buds of the vine during the plant’s annual growth cycle appears to be associated with different development phases (Pouget, 1963) and with abiotic stress as applied by the environment. In fact, the accumulation of those compounds during the phase of dormancy could be linked to the decrease in temperature and shorter length of daylight. Such results have also been suggested by many researchers (Macheix et al., 2005; Wahid and Ghazanfar, 2006) who have shown that certain polyphenolics also play a role in the plant’s tolerance to abiotic stress and argue that the temperature, as a polyphenolic metabolism expression regulator, induces an accumulation of anthocyanins. As such, this regulation may intervene at the level of activity Phenylalanine (PAL) (Macheix et al., 1990).

The accumulation of polyphenolic compounds seems to be associated with low temperatures, characterizing the phase of eco-dormancy when temperatures stay below 10°C for a period of 10 days (Pouget, 1963). Such a hypothesis seems in agreement with the findings of Pennycooke et al. (2005) who showed that the cold caused an accumulation of polyphenolic compounds in *Betunia hybrida* leaves. Wróbel et al. (2005) have also shown that the cold stimulates some polyphenolic compounds (free gallic acid and catechin) in the seeds of *Vitis riparia*.

According to the authors cited above, once those polyphenolic compounds metabolize, they are transferred to other parts of the plant to meet the plant’s developmental requirements. Such results were also yielded by Zohra et al. (2011) who reported that levels of quercetin in the grapevine variety Carignan increase during the phase of dormancy and decrease during the phase of budbreak. Be that as it may, the results of the present experiment did not confirm those findings since, in the case of the present study, quercetin increased during eco-dormancy and spiked once the treatments were effected (Table 4).

As mentioned earlier, the variations in polyphenolic compounds of latent buds during their annual growth cycle seems to be related to hydrogen peroxide (Li et al., 2011), but the combination of a cinnamic acid pretreatment and a cold treatment further enhances the antioxidant activities. Proline and total polyphenolics content increased under chilling stress (Shima et. al., 2017). Acclimation to low temperature has been shown to increase concentrations of polyphenolic compounds in several fruit tree species including the grapevine (Said et al., 2014; Karimi and Ershadi, 2015) and pistachio (Pakkish et al., 2009).

All authors cited above showed that, in association with cinnamic acid, cold increases H_2_O_2_ levels. That result seems to be in agreement with several studies dealing with the involvement of the H_2_O_2_ in response to attacks of the abiotic environment (Ashraf and Harris, 2004): it is quite likely that H_2_O_2_ acts as a messenger triggering a modification of ionic flux or a production of secondary messengers such as salicylic acid.

### Principal Component Analysis (PCA)

PCA transforms the original data set, all measurements included, into a smaller set of uncorrelated new variables (Principal Components, where eigenvalues are >1). When performed on the polyphenolics studied, the PCA produced two (2) components, in declining order of importance, which accounted for and explained 63.95% of the total variability between and among the different polyphenolics (Table 5).

**Table 5:** Principal components (PC) of the polyphenolics evaluated

| Component |  | Percent of variance | Cumulative percentage |
| --- | --- | --- | --- |
| Number | Eigenvalue |  |  |
| 1 | 6.23 | 48.24 | 48.24 |
| 2 | 2.01 | 15.71 | 63.95 |

Polyphenolics compounds are presented in Figure 1 as a function of both the first and second principal components (PC). The first principal component (PC1) accounted for 48.24% of the total variability and was defined by polyphenolic compounds catechin, chlorogenic acid, epicatechin, o-coumaric acid, piceid, procyanidin B1, procyanidin B2, rutin, vanillic acid, and ε-viniferin. The fact that all compounds were located at a distance from the axis origin, suggests that they were well represented by PC1 and were placed close together on the positive side of the PC1 to indicate a strong, positive correlation. The second principal component (PC2) explained another 15.71% of the total variability and was defined by luteolin, quercetin, and resveratrol which were also placed close together on the positive side of PC2, to indicate that they positively correlated with one another. In contrast, luteolin did not correlate with ε-viniferin, catechin, nor did it do so with any of the remaining polyphenols.

**Figure 1:**
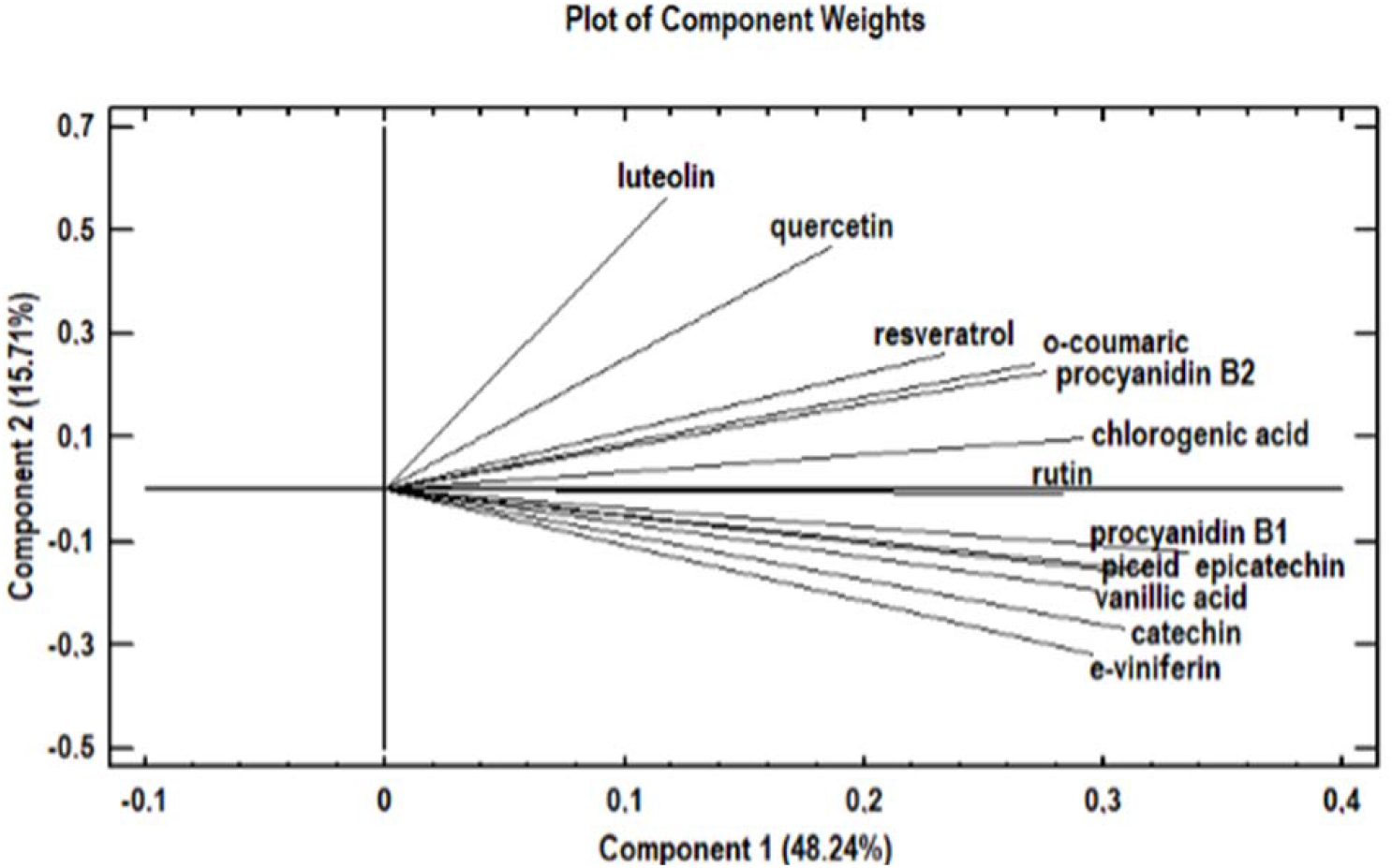
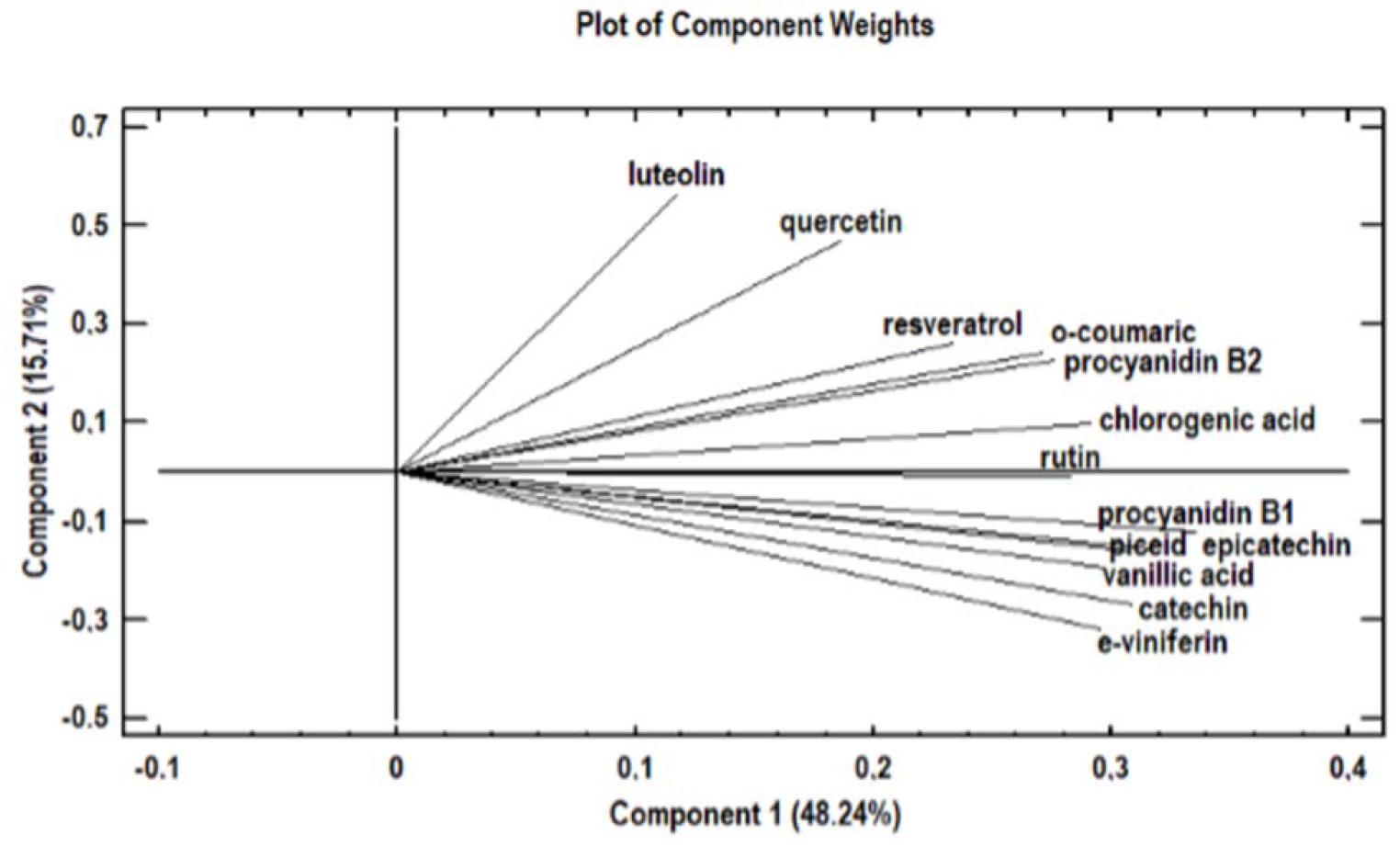
Principal Component Analysis: plot of the polyphenols in relation to the first two principal components (PC1 & PC2).

The first component contains the measurements gauging catechin, chlorogenic acid, epicatechin, o-coumaric acid, piceid, procyanidin B1, procyanidin B2, rutin, vanillicacid, and ε-viniferin. As to the second component, it gauges quercetin, resveratrol, and luteolin. According to the PCA analysis for the purposes of the present study, luteolin increased in contrast to catechin and ε-viniferin, as shown in Figure 1.

### Conclusions

The results of the present study showed that chemically-induced stress conditions, such as the ones caused by the application of chemical regulators which affect bud dormancy release and advance budbreak, also affect the content of polyphenolic compounds. Moreover, based on the experimental design that was implemented through (a) the different applications taking place on specific days and (b) the cane sampling taking place after a specific number of days had elapsed, the results confirmed, for the first time in such studies, that variations in the levels of the polyphenolic compounds identified do exist. Taking into consideration the fact that polyphenolic compounds are essentially involved in the resistance response to chilling stress, the variations in their levels constitute a first indicator of how much stress the latent buds actually undergo.

## Parsed Citations

Amiot MJ, Aubert S, Nicolas J (1993) Phenolic composition and browning susceptibility of various apple and pear cultivars at maturity and postharvest. Acta Hortic 343: 67–69 Google Scholar: Author Only Title Only Author and Title

Ashraf M, Harris PJC (2004) Potential biochemical indicators of salinity tolerance in plants. Plant Sci 166: 3–16 Google Scholar: Author Only Title Only Author and Title

Ayaz FA, Kadioglu A, Reunanen M (1997) Changes in phenolic acid contents of Diospyros lotus L. during fruit development. J Agric Food Chem 45: 2539–2541 Google Scholar: Author Only Title Only Author and Title

Babic I, Amiot MJ, Nguyen-The C, Aubert S (1993) Changes in phenolic content in fresh ready-to-use shredded carrots during storage. J Food Sci 58: 351–356 Google Scholar: Author Only Title Only Author and Title

Balasundram N, Sundram K, Samman S (2007) Phenolic compounds in plants and agri-industrial by products: Antioxidant activity, occurrence, and potential uses. Food Chem 99: 191–203 Google Scholar: Author Only Title Only Author and Title

Biniari K, Xenaki M, Daskalakis I, Rusjan D, Bouza D, Stavrakaki M (2020) Polyphenolic compounds and antioxidants of skin and berry grapes of Greek Vitis vinifera cultivars in relation to climate conditions. Food Chem 307: 125518 Google Scholar: Author Only Title Only Author and Title

Blankenship SM, Richardson DG (1985) Changes in phenolic acids and external ethylene during long-term cold storage of pears. J Am Soc Hortic Sci 110: 336–339 Google Scholar: Author Only Title Only Author and Title

Botelho RV, Pavanello AP, Pires EJP, Terra MM, Muller MML (2007) Effects of chilling and garlic extract on bud dormancy release in Cabernet Sauvignon grapevine cuttings. Am J Enol Vitic 58: 402–404 Google Scholar: Author Only Title Only Author and Title

Bouchereau A Aziz A, Larher F, Martin-Tanguy J (1999) Polyamines and environmental challenges: recent development. Plant Sci 140: 103–125 Google Scholar: Author Only Title Only Author and Title

Carreno J, Faraj S, Martinez A (1999) The effects of hydrogen cyanamide on budburst and fruit maturity of Thompson Seedless grapevine. J Hortic Sci Biotech 74: 426–429 Google Scholar: Author Only Title Only Author and Title

Challice JC, Williams AH (1968) Phenolic compounds of genus Pyrus–II: A chemotaxonomic survey. Phytochemistry 7: 1781 Google Scholar: Author Only Title Only Author and Title

Champagnat P (1983) Quelques réflexions sur la dormance des bourgeons des végétaux ligneux. Physiol Vég 21: 607–618 Google Scholar: Author Only Title Only Author and Title

Daskalakis I, Biniari K (2019) A new measurement model to estimate the intensity of acrotony on the latent buds of grapevine canes (Vitis vinifera L.). Not Bot Horti Agrobo 47: 3 Google Scholar: Author Only Title Only Author and Title

Dokoozlian NK, Williams LE, Neja RA (1995) Chilling exposure and hydrogen cyanamide interact in breaking dormancy in grape buds. Hortic Sci 30: 1244–1247 Google Scholar: Author Only Title Only Author and Title

Febvre E (1981) Contribution à l’étude du développement de boutures de Saule (Salix babylonica L.) cultivées in vitro. Influence de différentes températures de culture; induction d’une dormance et conséquences morphogènes. Thése Doct. 3ème cycle. Univ. Clermont-Ferrand Google Scholar: Author Only Title Only Author and Title

Herrmann K (1989) Occurrence and content of hydroxycinnamic and hydroxybenzoic acid compounds in foods. Crit Rev Food Sci Nutr 28(4): 315–347 Google Scholar: Author Only Title Only Author and Title

Ho CT, Lee CY, Huang MT (1992) Phenolic compounds in food and their effects on health. I. Analysis, Occurrence, and Chemistry (ACS Symposium Series 506) Google Scholar: Author Only Title Only Author and Title

Iwasaki K (1980) Effect of bud scale removal, calcium cyanamide, GA3 and ethephon on budbreak of ‘Muscat of Alexandria’ grape (Vitis vinifera L.). J Jpn Soc Hortic Sci 48: 395–398. Google Scholar: Author Only Title Only Author and Title

Karimi R, Ershadi A (2015) Role of exogenous abscisic acid in adapting of ‘Sultana’ grapevine to low temperature stress. Acta Physiol Plant 37(8): 1-11 Google Scholar: Author Only Title Only Author and Title

Kubota N, Miyamuki M (1992) Breaking bud dormancy in grapevines with garlic paste. J Am Soc Hortic Sci 7(6): 898–901 Google Scholar: Author Only Title Only Author and Title

Kuroi I (1985) Effect of calcium cyanamide and cyanamide on budbreak of ‘Kyoho’ grape. J Jpn Soc Hortic Sci 54: 301–306 Google Scholar: Author Only Title Only Author and Title

Lavee S, May P (1997) Dormancy of grapevine buds-facts and speculation. Aust J Grape Wine Res 3: 31–46 Google Scholar: Author Only Title Only Author and Title

Li Q, Yu B, Gao Y, Dai AH, Bai JG (2011) Cinnamic acid pretreatment mitigates chilling stress of cucumber leaves through altering antioxidant enzyme activity. J Plant Physiol 168: 927–934 Google Scholar: Author Only Title Only Author and Title

Lin CH, Wang TY (1985) Enhancement of bud sprouting in grape single bud cuttings by cyanamide. Am J Enol Vitic 36: 15–17 Google Scholar: Author Only Title Only Author and Title

Macheix JJ, Fleuriet A, Billot J (1990) Fruit phenolics, CRC Press, Inc., Boca Raton, Florida, USA

Macheix JJ, Fleuriet A, Jay-Allemand C (2005) Les composés phénoliques des végétaux: un exemple de métabolites secondaires d’importance économique. Lausanne: Presses polytechniques universitaires romandes. Print. Google Scholar: Author Only Title Only Author and Title

Manach C, Scalbert A, Morand C, Rémésy C, Jiménez L (2004) Polyphenols: Food sources and bioavailability. Am J Clin Nutr 79: 727–747 Google Scholar: Author Only Title Only Author and Title

Mizutani F, Shinohara K, Amano S, Hino A Kadoya K, Akiyoshi H, Watanabe J (1994) Effect of KCN and SHAM on budbreak and rooting of single eye cuttings of ‘Kyoho’ grape. Bull Exp Farm Coll Agr Ehime Univ 15: 1–5 Google Scholar: Author Only Title Only Author and Title

Nagar PK, Sood S (1996) Changes in endogenous abscisic acid and phenols during winter dormancy in tea (Camellia sinensis L. O. Kuntze. Acta Physiol Plant 18: 33–38 Google Scholar: Author Only Title Only Author and Title

Nigond J (1967) Recherches sur la dormance des bourgeons de la vigne. Thése de Doctorat. Université de Paris Google Scholar: Author Only Title Only Author and Title

Nortje BK, Koeppen BH (1965) The flavonol glycosides in the fruit of Prunus communis L. cultivar Bon Chrétien. Biochem J 97(1): 209–213 Google Scholar: Author Only Title Only Author and Title

Oleszek W, Amiot MJ, Aubert SY (1994) Identification of some phenolics in pear fruit. J Agric Food Chem 42: 1261–1265 Google Scholar: Author Only Title Only Author and Title

Olsen JE (2006) Mechanisms of dormancy regulation. Acta Hortic 727: 157–165 Google Scholar: Author Only Title Only Author and Title

Pakkish Z, Rahemi M, Baghizadeh A (2009) Seasonal changes of peroxidase, polyphenol oxidase enzyme activity and phenol content during and after rest in pistachio (Pistacia vera L.) flower buds. World Appl Sci J 6(9): 1193–1199 Google Scholar: Author Only Title Only Author and Title

Peer WA, Brown DE, Tague BW, Muday GK, Taiz L, Murphy AS (2001) Flavonoid accumulation patterns of transparent testa mutants of arabidopsis. Plant Physiol 126(2): 536–548 Google Scholar: Author Only Title Only Author and Title

Pennycooke JC, Cox S, Stushnoff C (2005) Relationship of cold acclimation, total phenolic content and antioxidant capacity with chilling tolerance in petunia (Petunia × hybrida), Environ Exp Bot 53(2): 225–232

Pouget R (1963) Recherches physiologiques sur le repos végétatif de la vigne (Vitis vinifera L.): La dormance des bourgeons et le mécanisme de sa disparition. Thése Dr Sc. Nat., no 123, Bordeaux, pp. 245 Google Scholar: Author Only Title Only Author and Title

Said Q, Ez-zohra IF, Mohamed F, Tayeb K (2014) Changes in levels of hydrogen peroxide and phenolic compounds in grapevine latent buds during the annual cycle. Int J Sci Res Public 4(5): 1–5 Google Scholar: Author Only Title Only Author and Title

Shima R, Ghasem S, Mehdi G, Mansour G, Ehsan M, Rouhollah K (2017) Effect of gradual and shock chilling stress on abscisic acid, soluble sugars and antioxidant enzymes changes in ‘Sultana’ grapevine. Iran J Plant Physiol 7(4): 2211–2224 Google Scholar: Author Only Title Only Author and Title

Shulman Y, Nir G, Fanberstein L, Lavee S (1983) The effect of cyanamide on the release from dormancy of grapevine buds. Sci Hortic 19: 97–104 Google Scholar: Author Only Title Only Author and Title

Wahid A, Ghazanfar A (2006) Possible involvement of some secondary metabolites in salt tolerance of sugarcane. J Plant Physiol 163: 723–30 Google Scholar: Author Only Title Only Author and Title

Weaver RJ, McCune SB, Coombe BG (1961) Effects of various chemicals and treatments on rest period of grape buds. Am J Enol Vitic 12: 131–142 Google Scholar: Author Only Title Only Author and Title

Wróbel M, Karama M, Amarowicz R, Frczek E, Weidner S (2005) Metabolism of phenolic compounds in Vitis riparia seeds during stratification and during germination under optimal and low temperature stress conditions. Acta Physiol Plant 27(3): 313–320 Google Scholar: Author Only Title Only Author and Title

Zelleke A, Kliewer WM (1989) The effects of hydrogen cyanamide on enhancing the time and amount of budbreak in young grape vineyards. Am J Enol Vitic 40: 47–51 Google Scholar: Author Only Title Only Author and Title

Zohra KD, Asma Z, Kamel M, Helmi H, Béchir E (2011) Changes of phenolic compounds in Carignan merithallus (Vitis vinifera L.) during bud dormancy and end of dormancy phase: correlation with rhizogenesis. Agric Sci 2: 498–504 Google Scholar: Author Only Title Only Author and Title

